# DUE-B Is Dispensable for Early Development and DNA Replication in Vertebrates

**DOI:** 10.1101/2025.06.17.658125

**Authors:** Courtney G. Sansam, Larissa A. Sambel, Emily A. Masser, Duane Goins, Gheorghe Chistol, Christopher L. Sansam

## Abstract

The DNA Unwinding Element-Binding protein (DUE-B) is a Cyclin Dependent Kinase (CDK) and Dbf4-Dependent Kinase (DDK) substrate that has been implicated in the control of DNA replication initiation. Previous studies reported that knocking down DUE-B in HeLa cells perturbs the G1-to-S phase transition, while depleting DUE-B from interphase *Xenopus* egg extracts impairs replication initiation. Based on these findings, the prevailing view is that DUE-B is a vertebrate-specific DNA replication initiation factor. Here, we asked whether *due-b* was an essential vertebrate gene *in vivo*, and whether it was critical for proper embryonic development in the zebrafish *Danio rerio*. We have generated *due-b* mutant zebrafish through genome-editing TALENs that fail to express *due-b* mRNA or protein. These mutant zebrafish are viable and survive to adulthood. They do not display outward developmental phenotypes, and when stressed with replication inhibitors, do not differ from their wild-type counterparts. Cell cycle analysis demonstrates that DNA replication occurs normally. Consistent with the zebrafish data, immunodepleting DUE-B from *Xenopus* nuclear egg extract did not impair DNA replication. Taken together, our findings indicate that DUE-B is dispensable for DNA replication and early development in vertebrates.

**Summary:** The DNA replication factor DUE-B is not required for zebrafish development or genome duplication, suggesting it plays a redundant or specialized role in DNA replication.

## Introduction

DNA replication is strictly orchestrated so that the entire genome is replicated accurately and to completion during each cell cycle, prior to the onset of mitosis. Dysregulation of DNA replication is associated with primordial dwarfism, neuro-developmental disorders, and genomic instability (1, 2). DNA replication is controlled at two distinct stages: origin licensing and origin firing. During origin licensing, ORC, CDC6, and CDT1 load two MCM2-7 hexamers at potential origins to form the <u>pre</u>-replicative <u>c</u>omplex (pre-RC) (3, 4). Origin firing entails the assembly and activation of two replicative DNA helicases at each origin, which requires several additional replication proteins called firing factors (5–12). In yeast, phosphorylation of MCM2-7 by the <u>D</u>bf4-<u>d</u>ependent protein <u>k</u>inase (DDK) stimulates the recruitment of firing factors Sld3 and Sld7 to select licensed origins (13, 14). S-phase specific phosphorylation of Sld2 and Sld3 by <u>C</u>yclin <u>D</u>ependent <u>K</u>inase (S-CDK) facilitate the recruitment of GINS and CDC45 subunits to the pre-RC, resulting in the assembly of the replicative helicase CMG (CDC45•MCM2-7•GINS) (12, 15). Following its activation in a MCM10-dependent manner, the CMG helicase begins unwinding DNA, thus initiating DNA replication (16).

Vertebrate origin licensing and firing is broadly similar to yeast with some notable differences. For instance, vertebrate TRESLIN (also known as TICRR) is functionally analogous to Sld3, but differs significantly in both structure and size, with only remote protein homology to the yeast Sld3 (5, 6, 17). Metazoan MTBP is triple the size of its yeast ortholog Sld7, although it still is an obligate binding partner to TRESLIN/Sld3 (7, 18, 19). Similarly, metazoan RECQL4 is ∼3x larger than its yeast ortholog Sld2, contains metazoan-specific domains, and has poor sequence homology to Sld2 (20, 21). In contrast to these poorly conserved initiation factors, many other core replication proteins (including CMG subunits CDC45, MCM2-7, and GINS) are highly conserved between yeast and vertebrates. Compared to yeast, metazoa have much larger genomes and a more complex chromatin environment, likely requiring additional regulators of replication initiation (6, 18). For example, DONSON was recently identified as a novel essential origin firing factor in higher eukaryotes with no direct ortholog in yeast (22–24). This raises the question if there are other essential regulators of replication initiation that are unique to metazoa. Over the years, a few understudied proteins have been implicated in regulating origin firing in vertebrates, notably GEMC1, HELB, and DUE-B (also known as <u>D</u>-aminoacyl-tRNA <u>d</u>eacylase 1, or DTD1) (25–31).

<u>D</u>NA <u>u</u>nwinding <u>e</u>lements (DUE) were identified in *E.coli* and in yeast as thermodynamically unstable sequences capable of helical unwinding (32). These unique sequence sites were proposed to serve as sites of replication initiation. The <u>D</u>NA <u>U</u>nwinding <u>E</u>lement-<u>B</u>inding protein (DUE-B) was originally identified in a yeast one-hybrid screen for proteins affecting DNA replication initiation at the c-Myc origin site (8). This well-characterized replication site contains a known replication initiation zone and a DNA Unwinding Element that supports both transcription factor and replication factor binding (8, 33–36). Deletion of the c-Myc DUE region or deletion of DUE-B protein inhibited its origin activity (37). Chromatin-immunoprecipitation in HeLa cells confirmed the presence of DUE-B protein on chromatin at the c-Myc origin site, as well as at the Lamin-B2 core replicator origin site (8, 36).

Yeast DUE-B is a 209 amino acid protein and has been classified as a non-essential D-aminoacyl-tRNA deacylase (8, 38). Intriguingly, vertebrate DUE-B has acquired an additional 60 amino acid C-terminal tail that is not present in either *S. cerevisiae* or *C. elegans*. This C-terminal extension has been reported to interact with TRESLIN, TOPBP1, and CDC45 proteins in HeLa cells (30, 31). It was previously reported that in interphase *Xenopus* egg extracts DUE-B is required for CDC45 and TOPBP1 recruitment to chromatin (i.e. origin firing), but is not required for MCM2-7 recruitment to chromatin (i.e. origin licensing). Importantly, depleting DUE-B from interphase egg extract strongly impaired DNA replication, and this defect was rescued by supplementing depleted extract with recombinant DUE-B (31). Studies performed in HeLa cells reported that knocking down DUE-B delayed the G1-S transition and prevented TRESLIN and CDC45 binding to chromatin (8, 30). Based on these multiple lines of evidence, it was proposed that DUE-B is necessary for replication initiation in higher eukaryotes, strongly suggesting that DUE-B should be essential for proper development in vertebrates (31).

Thus far, studies of DUE-B in DNA replication have been limited to cell culture and cell-free systems. To determine whether DUE-B is essential during vertebrate embryonic development, we generated and analyzed *due-b* mutant zebrafish. Zebrafish are well suited for studying cell cycle and DNA replication genes during development, as their transparent, externally developing embryos are easily accessible for developmental analysis and genetic manipulation (39–41). Furthermore, maternally deposited mRNA and protein in zebrafish embryos often circumvent early developmental defects, allowing the functional assessment of essential genes at later embryonic or larval stages. Extensive characterization of zygotic mutants for DNA replication factors—including TICRR (TRESLIN), MCM5, GINS1/2, TopBP1, POLD1, and RIF1—has shown that most display defects in late-proliferating regions such as the eyes, brain, craniofacial structures, and hematopoietic tissues, while early development proceeds normally due to maternal mRNA and protein (5, 42–49). In rare cases where DNA replication mutants survive to adulthood, maternal-zygotic mutants (from crosses of homozygous mutant parents) can reveal more severe embryonic phenotypes due to the absence of both maternal and zygotic gene contributions (45).

We generated a TALEN-induced *due-b* mutant zebrafish carrying a frameshift mutation that results in severely reduced *due-b* mRNA levels and the complete loss of detectable protein expression. Unexpectedly, these mutants are viable and survive to adulthood in Mendelian ratios.Maternal-zygotic *due-b* mutants, which lack maternally supplied *due-b* mRNA and protein, also survive and show no early developmental delays or defects. They exhibit no overt developmental phenotypes or cell division defects during embryogenesis, even when challenged with replication inhibitors. Cell cycle analysis and EdU incorporation confirm normal DNA replication, and early cleavage divisions proceed with normal timing. Although mutants display a modest reduction in eye size, they retain the ability to regenerate caudal fins, indicating that *due-b* is dispensable for DNA replication in both embryonic and adult proliferative contexts. To further assess whether DUE-B is required for DNA replication in vertebrate embryos, we used nucleoplasmic *Xenopus* extracts that replicate DNA without the need to assemble a nuclear envelope (50). In our extract-based biochemical assays, immunodepleting the DUE-B protein did not impair DNA replication. Together, these results demonstrate that *due-b* is not an essential gene for genome duplication or for metazoan development.

## Results

### TALEN induced frameshift mutation in *due-b* causes a sharp decrease in mRNA

The C-terminal tail of the vertebrate DUE-B protein interacts with TRESLIN, TOPBP1, and CDC45 and was reported to be necessary for DNA replication in interphase *Xenopus* egg extracts (31). To assess the degree of conservation of the DUE-B C-terminal tail across species, we aligned the C-terminal protein sequences of DUE-B from yeast to humans (Fig. 1A). Vertebrates, including zebrafish and frogs, exhibited a high degree of identity with the human DUE-B protein. Specifically, the zebrafish *Danio rerio* DUE-B C-terminal sequence shares 76% identity with the human version, while *Xenopus laevis* exhibits 78% identity.

**Figure 1:**
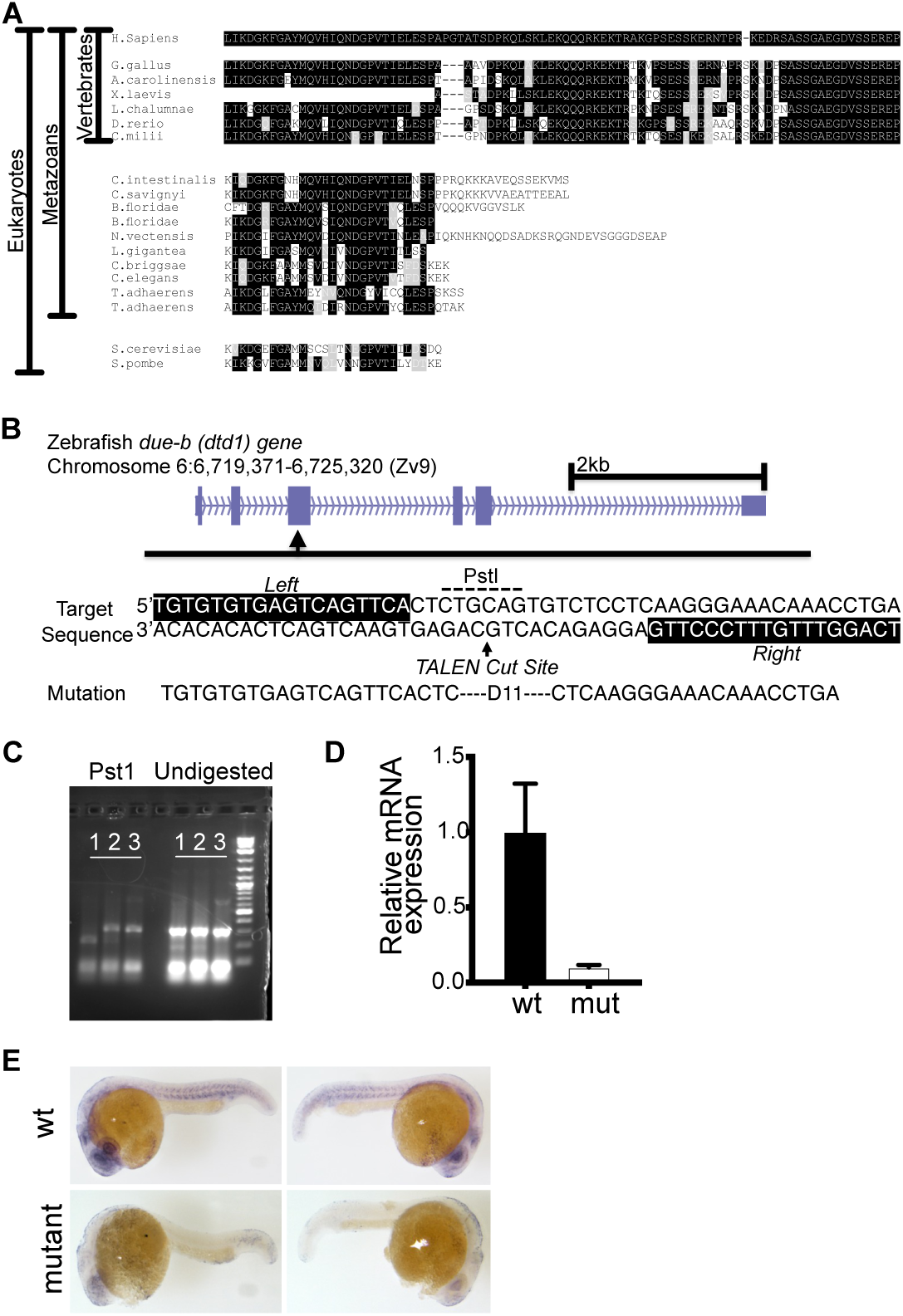
TALEN induced frameshift mutation in due-b causes a sharp decrease in mRNA. **(A)** DUE-B C-terminal protein across species. The C-terminal extension is highly conserved but only found in vertebrates. **(B)** Schematic of TALEN designed to cut the *due-b* (also known as *dtd1*) gene on Chromosome 6 at an existing PstI restriction site. **(C)** We identified a Founder fish carrying an 11bp deletion. PCR amplification and digestion with PstI. Lanes show wt, het, and mutant in order. **(D)** Quantitative PCR shows sharp reduction in mRNA levels in mutant fish. **(E)** *In situ* hybridizations for *due-b* in wild-type and *due-b* mutant fish. *due-b* is expressed in the developing head, eye and central nervous system, while mutant fish display strong reduction in signal.

Given the high conservation of the zebrafish DUE-B sequence, we devised a strategy to generate a zebrafish mutant lacking the DUE-B protein. Zebrafish has a single copy of the *due-b* gene (also known as *dtd1*) on Chromosome 6. Using a transcription activator-like effector nuclease (TALEN) known for precise genome editing, we designed a targeted cut within exon 2 of *due-b* focusing on a specific PstI restriction site (Fig 1B) (39). We injected *due-b* site-specific TALENs into one-cell stage wild-type Tubingen AB5 (TAB5) zebrafish embryos, raised them to adulthood, mated them with wild-type TAB5 fish, and screened their F1 offspring for mutations. Through PCR amplification around the TALEN cut site and subsequent PstI digestion, we identified fish resistant to digestion, signifying a mutation at the restriction site sequence (Fig 1C, D). Sequencing confirmed that a zebrafish (F1-22) carried an 11bp deletion in *due-b*, leading to a frameshift mutation (Fig.1B). Typically, frameshift mutations in early exons trigger mRNA degradation via non-sense mediated decay, resulting in a deficiency of the encoded protein.

Confirmed F1-22 heterozygotes were bred to generate *due-b^+/+^, due-b^+/Δ11^*, and *due-b* ^Δ11/Δ11^ clutch mates (hereafter referred to as *due-b^omf202/omf202^*). To assess *due-b* mRNA expression in these mutants, we genotyped individual embryos at 24 hours post fertilization (hpf), pooled like genotypes, and quantified mRNA levels using real-time PCR (qRT-PCR). Zebrafish embryos identified as *due-b^omf202/omf202^*exhibited a significant reduction in *due-b* mRNA expression (Fig 1D). Next, we evaluated the extent of mRNA depletion throughout the whole embryo by performing *in situ* hybridizations of *due-b* in 24 hpf embryos. The *in-situ* results show that *due-b* mRNA is highly expressed in the eye, brain, and along the developing central nervous system, all sites of rapid cell division at 24 hpf embryos (Figure 1E). The *due-b^omf202/omf202^* embryos lacked detectable mRNA expression at these sites (Figure 1E).

### *due-b^omf202/omf202^* zygotic mutants are viable

Incross matings of heterozygous zebrafish carrying mutations in components of the pre-RC such as *mcm3*, or firing factors such as *ticrr(treslin)* and *gins,* are not viable past 5 days post fertilization (dpf), and the embryos show severe developmental deformities, including an underdeveloped head, severe spinal curvature, and small eyes (5, 43). From our original heterozygous incross, we observed that the *due-b^omf202/omf202^* mutant embryos survived through 24 hpf, but we questioned whether they displayed any delays in development. Heterozygote F1-22 zebrafish were mated to produce wild-type, heterozygous and mutant clutchmates, and their progeny raised to evaluate long-term survival and developmental growth. We carefully monitored *due-b^omf202/omf202^*clutches for survival and developmental deformities throughout the first few days of development (Fig. 2A). Unexpectedly, *due-b^omf202/omf202^* embryos were viable, showed no overt developmental phenotype at 24 hpf, and were free-feeding by 5 dpf. The *due-b^omf202/omf202^*zebrafish reached sexual maturity at expected Mendelian ratios (Fig 2B). By Day 70, we measured the length of each fish to determine if *due-b^omf202/omf202^* zebrafish matured with an overall smaller size. While wild-type and heterozygous fish had mean lengths of 2.05cm (*n=47*) and 2.13cm *(n=100)* respectively, *due-b^omf202/omf202^* zebrafish were similarly sized, with a mean length of 2.19cm *(n=52)*. The differences in overall length did not reach statistical significance based on genotype (One-way ANOVA, Fig 2C). To confirm whether DUE-B protein was absent in these mutants, we collected synchronized wild-type, heterozygous, or *due-b^omf202/omf202^* embryos at 10 hpf. We dechorionated and deyolked the embryos, disaggregated the cells, and made whole cell protein lysate using RIPA buffer and immunoblotted for DUE-B (antibody gift of M. Leffack). While a band of 24 kDa was detected readily in the wild-type sample, the *due-b^omf202/omf202^* embryos lacked this band, confirming loss of *due-b* protein product (Figure 2D).

**Figure 2:**
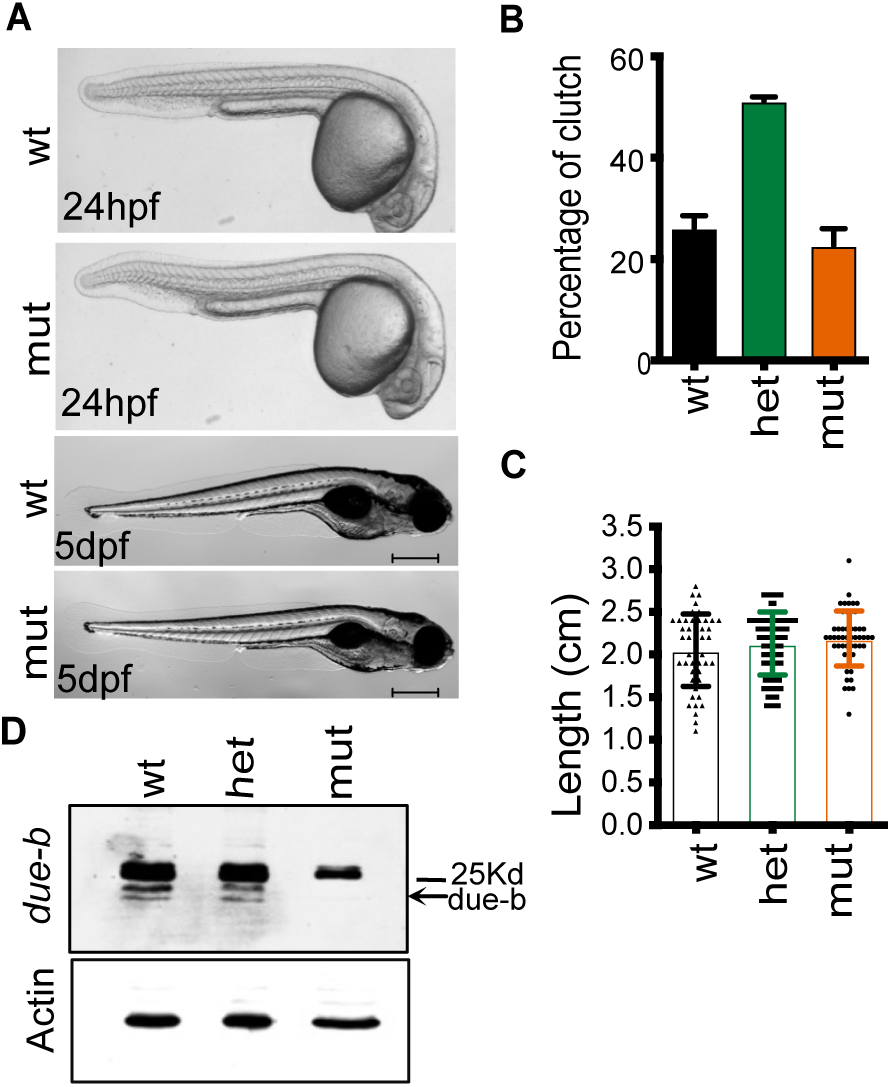
due-b^omf202/omf202^ zygotic mutants are viable. **(A)** Wild-type and *due-b*^omf202/omf202^ embryos at 24 hpf and 5 dfp do not show developmental differences. **(B)** Heterozygous *due-b*^+/omf202^ was incrossed to generate matched clutch mates. Normal Mendelian distributions were observed (n= 5 separate crosses). **(C)** Individual clutch mates from a Het incross were measured for length at Day 70 and data was evaluated using a One-Way ANOVA. **(D)** Western Blot for *due-b* protein in 10 hpf embryos. B-actin loading control.

### Maternal loss of DUE-B does not affect development

We were concerned that the lack of overt phenotype observed in the *due-b^omf202/omf202^* embryos could be a consequence of maternal mRNA contributions supplied by the mother in the first stages of development. Unlike mammals, zebrafish and frog embryos develop outside the mother and do not commence zygotic transcription until the mid-blastula transition (MBT) at approximately the 1000 cell stage (3 hpf). Thus, embryos can survive off maternally deposited mRNAs in the yolk for up to 5 dpf, until maternal mRNA stores are completely depleted. To test whether maternal *due-b* mRNA masks early developmental defects in the *due-b^omf202/omf202^*embryos, we mated adult homozygous *due-b^omf202/omf202^* zebrafish to generate maternal-zygotic mutants, which lack all maternal contribution of the *due-b* mRNA. To confirm efficient depletion of *due-b* transcripts in these embryos, we collected synchronized wild-type and *due-b^omf202/omf202^* embryos at 6 hpf (shield stage) and performed quantitative RT-PCR. As observed in the zygotic mutants, *due-b* mRNA levels were nearly undetectable in maternal-zygotic embryos compared to wild-type controls (1 ± SD 0.117 vs. 0.04 ± SD 0.0084) (Fig. 3A). We also assessed protein expression at 10 hpf and again found that the 24 kDa DUE-B protein was readily detected in wild-type lysates but absent in the mutant embryos (Fig. 3B).

**Figure 3:**
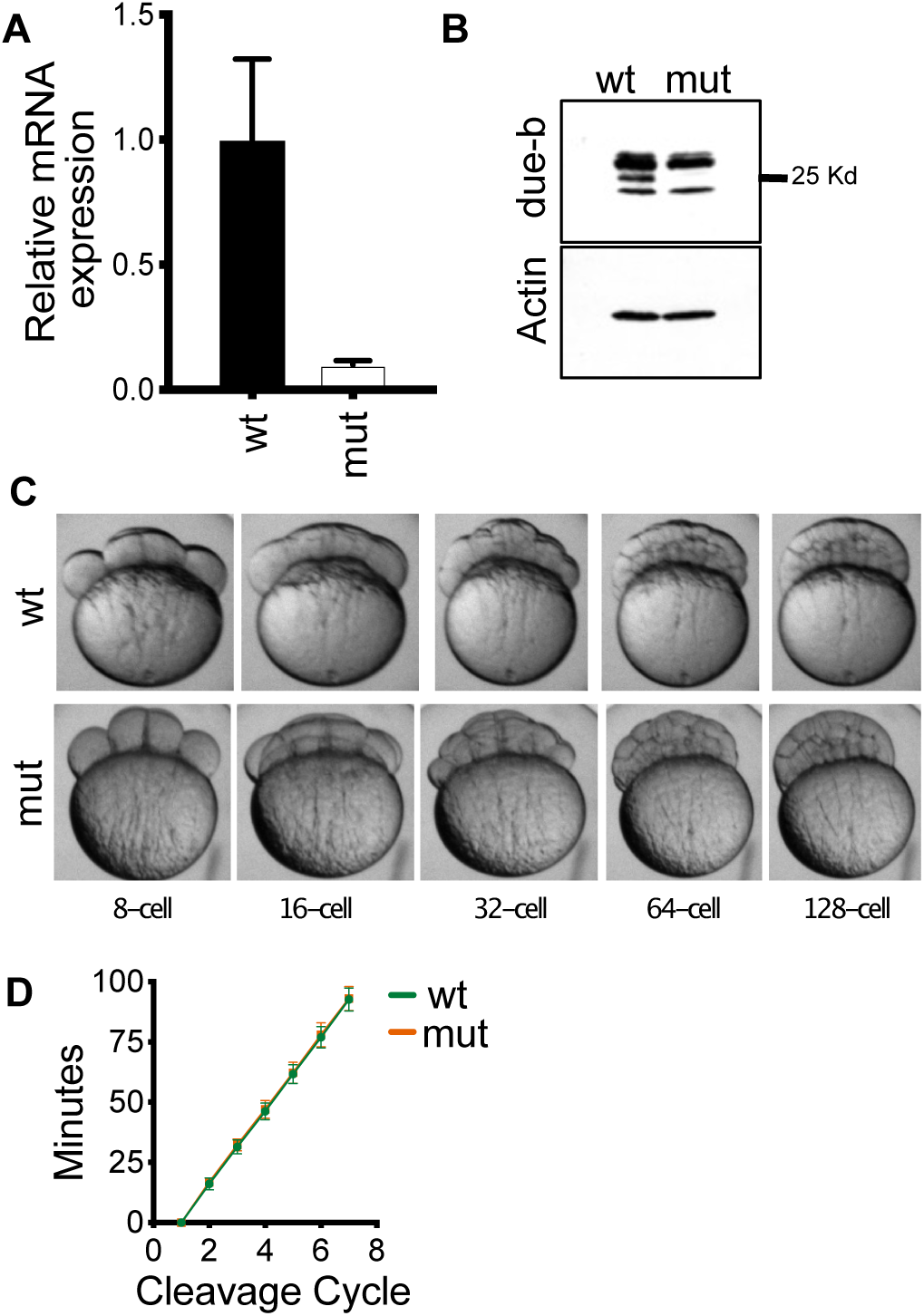
Maternal loss of due-b does not affect early development. **(A)** Quantification of total mRNA expression shows that the levels of *due-b* mRNA are greatly reduced in *due-b* maternal zygotic mutants. **(B)** Western Blot for DUE-B protein in 10 hpf wild type or maternal zygotic mutant embryos. B-actin loading control. (**C, D)** Synchronized wild-type or mutant embryos were imaged for the first eight cell divisions and the timing of cleavage events was recorded. No differences in cell division time was observed (n=10 wt embryos, n=11 mutant embryos).

Zebrafish embryos, from fertilization at 0 hpf to 3 hpf, are remarkable for their synchronous, rapid cleavage cycles. These early cleavage events represent continuous cycles of DNA replication followed by mitosis without gap phases (G1 and G2). Cleavages occur every 15 minutes until the cell cycle lengthens following the 10^th^ cell division at the Mid-blastula transition (MBT). The lengthening of S phase at this timepoint is accompanied by a loss of synchrony in cell divisions (51). Since depleting DUE-B was reported to impair the loading of Cdc45 onto chromatin in both *Xenopus* extracts and in HeLa cells (8, 29–31, 36, 37), we reasoned that DUE-B may be critical for these early rounds of DNA replication. We considered the Cleavage Cycle possibility that the early cell cycles were compromised in our *due-b^omf202/omf202^* embryos, but that the embryos ultimately recovered and survived to adulthood at expected ratios. Detecting such defects are a particular strength of the zebrafish system. We therefore monitored these early cleavage cycles in wild type and *due-b^omf202/omf202^* maternal-zygotic mutant embryos by time-lapse photography (Fig. 3C). We timed the cell cleavage events in each embryo and plotted these division cycles against time (Fig. 3D). Our results show identical cleavage cycle times for wild-type and *due-b* mutant embryos indicating that loss of the DUE-B protein does not impair these early developmental cycles.

### DNA replication is normal in *due-b* mutant embryos

To directly assess DNA replication in the *due-b* mutants, we evaluated their ability to synthesize DNA using the thymidine analog 5-ethynyl-2’-deoxyuridine (EdU). At 24 hpf, wild-type, heterozygous, and *due-b* mutants were incubated in EdU for 20 minutes, after which EdU incorporation was quantified by flow cytometry (Fig. 4A). Surprisingly, embryos deficient for *due-b* showed no impairment in their ability to incorporate EdU. Furthermore, the distribution of cells across the different phases of the cell cycle was similar among all genotypes, indicating normal DNA replication and cell cycle progression in *due-b* mutant zebrafish (Fig. 4B).

**Figure 4:**
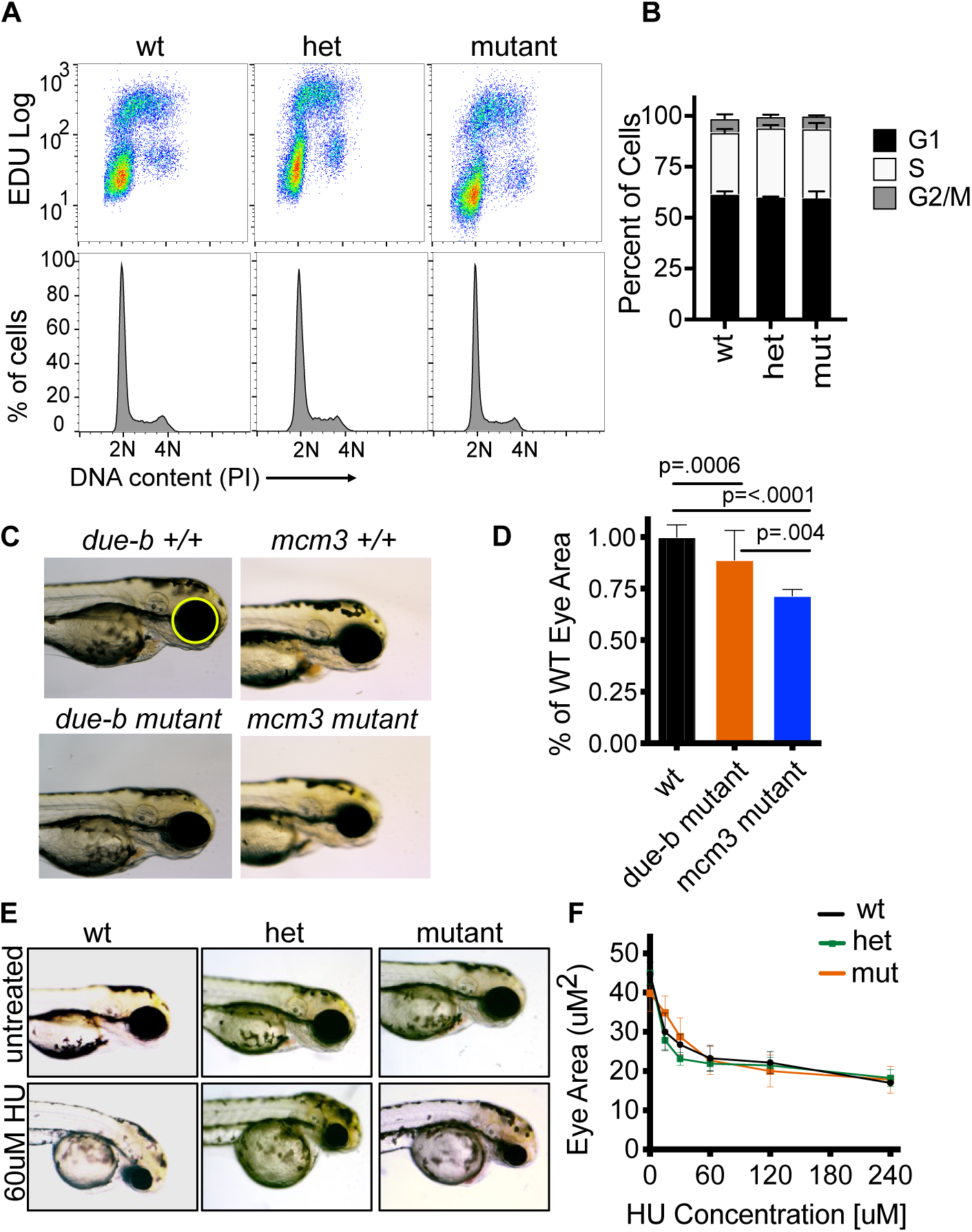
Marginal DNA replication defects in due-b mutants. **(A)** EdU incorporation and propidium iodide (PI) flow cytometry analysis of cells from pools of EdU pulse-labeled 24 hpf wild-type, heterozygous, and *due-b* mutant embryos (n=20,000 cells evaluated). **(B)** Quantification of flow cytometry data. Cells with G1, S, or G2/M DNA content do not show any difference in cell cycle phases between genotypes. **(C**) Eye area images of 72 hpf wt and *due-b* mutant embryos (left) or wt and *mcm3* mutant embryos (right). **(D)** Quantification of eye area was performed using ImageJ. Eye area was normalized to wild-type fish eyes. Statistical analysis using One-Way ANOVA showed significant differences between wt and *mcm3* mutants (p < 0.0001), between wt and *due-b* mutant (p = 0.0006) as well as between *mcm3* mutants and *due-b* mutants (p = 0.0041). **(E,F)** 24 hpf embryos were placed in E3 fish water containing different concentrations of hydroxyurea, ranging from 0 uM to 240 uM. The eye area was imaged and quantified at 72 hpf. There was no difference in eye area between genotypes in response to hydroxyurea treatment.

### Developmental eye growth is largely preserved in *due-b* mutants

The developing zebrafish eye is one of the most prominent and rapidly growing organs during early embryogenesis, with extensive proliferation and tissue expansion occurring between 16 and 72 hpf (43). Because this growth depends on robust DNA replication, measurements of eye size can serve as a sensitive readout of proliferative capacity during development. We therefore asked whether *due-b* mutant embryos exhibit reduced eye size as an indicator of compromised replication. To address this, we imaged and measured the eye area of individual 72 hpf *due-b^omf202/omf202^* embryos and compared them to wild-type siblings (Fig. 4C-D). As a benchmark for replication-dependent eye growth, we also analyzed 72 hpf embryos from an *mcm3^^+/HI3068^* incross. MCM3 is a core component of the pre-RC and the replicative helicase CMG, and loss of *mcm3* has been shown to impair eye development in developing zebrafish embryos (43). Quantification revealed that *due-b* mutants had a modest but statistically significant reduction in eye area (49.1 µm, *n* = 45) compared to wild-type embryos (53.4 µm, *n* = 28; *p* = 0.0004, one-way ANOVA). By comparison, *mcm3* mutant embryos exhibited a more pronounced phenotype, with an average eye area of 43.7 µm (*n* = 7) (Fig 4D).

### DUE-B loss does not increase hydroxyurea sensitivity

Having shown little or no effect on proliferation during development in *due-b* mutant embryos, we next asked whether replication stress might uncover a more subtle role for DUE-B in DNA replication. Only a fraction (5-10%) of licensed replication origins are normally activated during S phase, while the remainder function as dormant origins that can be used when replication forks stall (52, 53). Hydroxyurea (HU) induces replication stress by inhibiting ribonucleotide reductase, leading to dNTP depletion and fork stalling, thereby increasing reliance on dormant origin activation (52, 54). Sensitivity to HU is a hallmark of mutations that impair replication initiation or fork stability (9, 12, 52, 54–57). To test whether DUE-B is required under these conditions, we treated wild-type and *due-b^omf202/omf202^*embryos with increasing concentrations of HU from 24 to 72 hpf and measured eye size as a readout of cellular proliferation during DNA replication stress. HU reduced eye size in both genotypes in a dose-dependent manner, but *due-b* mutants did not show increased sensitivity, suggesting that dormant origin usage remains largely intact in the absence of *due-b* (n = 8 per genotype; Fig. 4E-F).

### Adult zebrafish can regenerate the caudal fin in absence of DUE-B

We had initially focused on DNA replication in the development of the early embryo. We next considered the possibility that *due-b* may be essential for DNA replication in proliferating cells of adult tissues. We reasoned that a critical role of *due-b* could be in the re-initiation of DNA replication from quiescent, non-cycling cells. The zebrafish has the remarkable capability of regenerating many tissues, and regeneration of an amputated caudal tail fin in zebrafish is well characterized and easily monitored (58–60). Importantly, the zebrafish tail fin is comprised of terminally differentiated non-muscularized cells in the adult fish. To regenerate the tail following amputation, cells adjacent to the amputation site de-differentiate and cells of the blastema must re-enter the cell cycle and initiate rounds of DNA replication. Therefore, we asked if *due-b* played a role in regenerating tissue of the caudal fin. We amputated the caudal tail fins of wild-type, heterozygous, and *due-b* mutant (n=40), photographing each fish before and after surgery. We then single-housed the fish in tanks held at 33°C for 10 days. We photographed each fish on days 3, 5, 7, and 10 post-surgery and quantified the percent of tail regrowth (Fig. 5A-B). All the fish were able to regenerate their tail fins, and we found no significant difference in the rate of tail regeneration between wild-type and *due-b* mutant fish. This data indicates that DUE-B is not required for DNA replication re-initiation from non-cycling cell populations.

**Figure 5:**
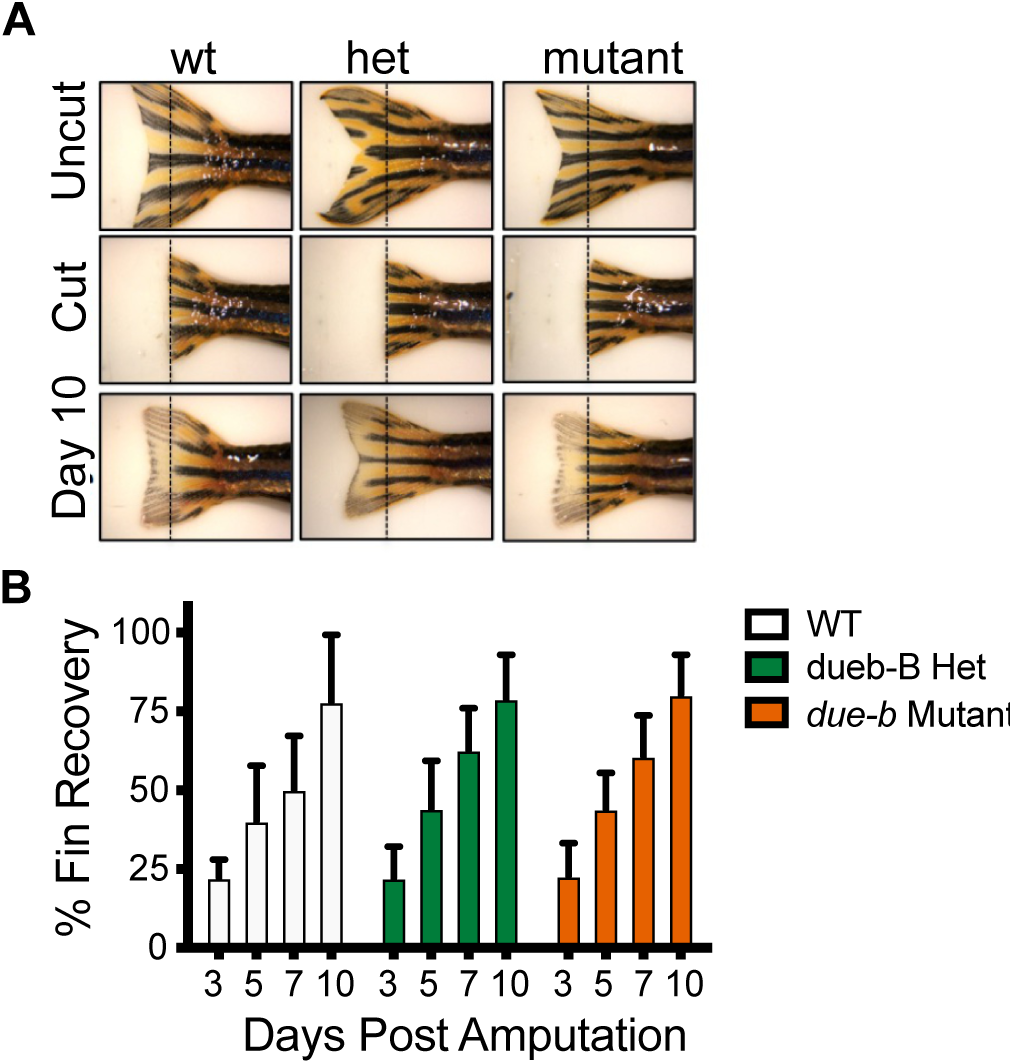
Adult zebrafish are able to regenerate caudal fin in absence of due-b. **(A)** The caudal fins of adult zebrafish were photographed, cut, and photographed again at day 0 (post-clip) 3, 7, and 10 days. Fin regrowth was measured as the number of pixels/mm^2^ of a line from the cut position to edge of new growth using Image J. **(B)** The percent of re-growth was calculated from the difference for wt (n=7), heterozygous (n= 18) and mutant fish (n=12).

### DUE-B is not required for DNA replication in *Xenopus* nuclear egg extract

In zebrafish, genetic compensation is a known phenomenon that can mask developmental phenotypes (61). Genetic compensation can occur through changes in transcriptional networks or pathway-level responses to gene disruption that result in enhanced viability. We reasoned that it was possible that the replication function of DUE-B was being masked in this way. Therefore, we sought to address the DNA replication function of DUE-B more directly in *Xenopus* egg extracts.

*Xenopus* egg extracts are a cell-free model for studying genome maintenance *in vitro,* and has been instrumental for understanding the molecular mechanisms of DNA replication (62). Notably, there are different versions of *Xenopus* extract, including cycling egg extract (63), interphase egg extract (64), and nuclear egg extract (50). DUE-B was previously implicated in DNA replication using interphase egg extract that can replicate sperm chromatin (31), but requires the formation of a nuclear envelope around chromatin to concentrate replication factors (64). As a result, perturbations that disrupt nuclear envelope function can impair DNA replication (65).

To investigate whether DUE-B is specifically required for DNA replication, we used *Xenopus* nuclear egg extract that can replicate a variety of DNA substrates in the absence of a nuclear envelope (50). We employed a plasmid replication assay (Fig. 6A) in which the template DNA is first licensed in High-Speed Supernatant (HSS) extract that mimics the G1 phase of the cell cycle, then replication is initiated by adding Nucleo-Plasmic Extract (NPE) that mimics S phase (66, 67).

**Figure 6:**
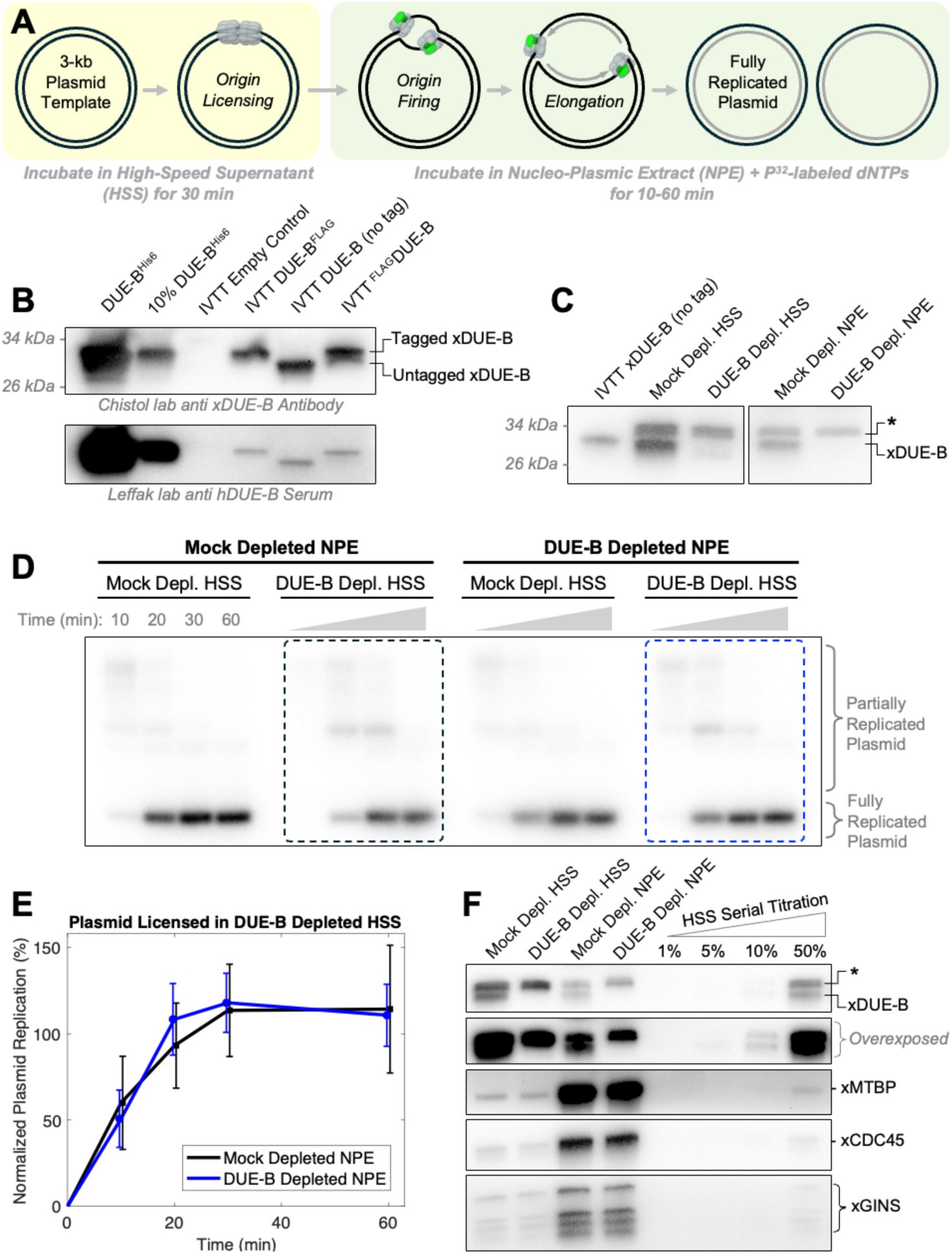
DUE-B is not required for DNA replication in Xenopus nuclear egg extract. **(A)** Diagram illustrating the DNA replication assay in *Xenopus* egg extract. **(B)** Immunoblots of recombinant DUE-B expressed in bacteria and in wheat-germ IVTT system. **(C)** Immunoblot of HSS and NPE egg extracts depleted with mock antibodies or anti-DUE-B antibodies (both images were cropped from the same blot image). **(D)** Autoradiograph of a replication gel with mock-depleted or DUE-B-depleted HSS or NPE. **(E)** Quantification of five independent repeats of the DNA replication assay. Error-bars represent the standard error. **(F)** Immunoblot of extracts used in the replication assay shown in panel (D).

To immunodeplete DUE-B from egg extract, we raised polyclonal rabbit antibodies against the full-length *Xenopus* DUE-B protein with a C-terminal His6-tag. Our affinity-purified anti-DUE-B antibodies efficiently detected recombinant *Xenopus* DUE-B purified from bacteria, as well as untagged *Xenopus* DUE-B expressed using Promega’s wheat-germ in vitro transcription-translation (IVTT) system (Fig. 6B). Polyclonal antibodies raised against full-length human DUE-B (a generous gift of Dr. Leffak) detected the same exact protein bands on a western blot (Fig. 6B). Our DUE-B antibodies detected two closely-spaced bands in both HSS and NPE extracts, but selectively immunodepleted only the lower band, which co-migrates with untagged *Xenopus* DUE-B expressed in IVTT (Fig. 6C). This led us to conclude that the lower band represents endogenous frog DUE-B in both egg extracts.

We replicated a 3-kb plasmid DNA template using *Xenopus* egg extracts supplemented with P^32^-labeled dCTP, which enabled us to visualize nascent DNA in a native agarose gel (Fig. 6D). DUE-B was not previously implicated in origin licensing (31) and is not required to reconstitute pre-RC loading with vertebrate proteins (4). For this reason, we licensed DNA in DUE-B-depleted HSS, and compared its replication in mock-depleted NPE versus DUE-B-depleted NPE (Fig. 6D, compare lanes in black box vs blue box). Immunodepleting DUE-B did not significantly alter the efficiency or kinetics of DNA replication (Fig. 6E), indicating that DUE-B is not required for DNA replication in *Xenopus* egg extracts without a nuclear envelope. Interestingly, unlike core replication initiation proteins (MTBP, CDC45, GINS) that are more abundant in the S-like extract (NPE), DUE-B is more abundant in the G1-like extract (HSS) (Fig. 6F)

## Discussion

We targeted *due-b* for mutagenesis in zebrafish because previous work in human cell culture systems and *Xenopus* egg extracts suggested that DUE-B is required for DNA replication initiation. Acute depletion studies reported delayed G1–S progression, defective CDC45 loading, and dependence on the conserved C-terminal extension for replication function (8, 29–31, 37, 38). These observations, together with DUE-B’s reported interactions with MCM2-7, CDC45, TRESLIN, and TOPBP1 (30, 31), led us to expect that *due-b* loss would disrupt replication and development *in vivo*.

Zebrafish assays, however, revealed that DUE-B is not required for genome duplication or normal development. *due-b* mutants are viable, achieve normal adult size, and progress through S phase without detectable delay. Their mild reduction in eye size contrasts sharply with the pronounced microphthalmia, body curvature, and early lethality observed in replication mutants such as *mcm3^HI3068^* and *ticrr^HI1573^* (5, 43). Furthermore, despite the high sensitivity of eye development to hydroxyurea, *due-b* mutants did not show increased susceptibility to replication stress. Finally, normal adult caudal fin regeneration in *due-b* mutants, a process that requires rapid and efficient proliferation of blastema cells, further supports the conclusion that DUE-B is dispensable for DNA replication in both embryos and adults. Together, these zebrafish studies indicate that loss of DUE-B does not compromise genome duplication under the developmental or stress conditions examined.

To complement these *in vivo* results, we interrogated DUE-B’s putative role in origin firing using biochemical replication assays in nuclear extracts prepared from *Xenopus* eggs. Previous reports described pronounced replication defects following acute DUE-B depletion in interphase *Xenopus* extracts, but our experiments did not reproduce these findings. This agreement between zebrafish mutants and *Xenopus* egg extracts suggests that DUE-B is not an essential replication initiation factor. Instead, DUE-B may act redundantly or in a condition-specific manner.

Although our data diverges from the results of earlier studies (8, 29–31, 36, 37), our work is supported by more recent genome-wide CRISPR screens showing that human DUE-B knockout has minimal fitness consequences compared to other essential replication genes (68, 69). DUE-B may therefore be required selectively, such as under oncogene-induced replication stress, when origin usage becomes dysregulated and additional stabilizing or activating factors may be needed. If DUE-B functions at specific origins or chromatin states, its loss might not affect bulk replication, but it could influence origin choice or timing in ways that remain undetected in our assays. Factors such as RECQL4 exhibit preferential control of baseline versus dormant origins (70), and DUE-B may similarly regulate a subset of initiation events.

One potential explanation for the robust fitness of the *due-b* mutant zebrafish is genetic or functional compensation. Zebrafish readily upregulate related genes in response to stable mutations, and we did not assess whether other known initiation factors such as TRESLIN (TICRR), MTBP, or DONSON become upregulated in *due-b* mutants. Compensation could similarly contribute to the lack of phenotype in human CRISPR knockout screens.

A potential explanation for the discrepancy between our experiments in DUE-B depleted nuclear *Xenopus* egg extract and previous experiments in DUE-B depleted interphase *Xenopus* egg extracts is the fact that DNA replication in interphase extract requires the formation of a nuclear envelope. It is possible that depleting DUE-B from interphase extract impaired nuclear envelope formation or transport, resulting in strong replication defect that could be rescued by adding back recombinant DUE-B. Interestingly, GEMC1 – another protein implicated in replication initiation, was also shown to be required for DNA replication in *Xenopus* interphase egg extracts (25). However, a more recent study showed that GEMC1-knockout mice are viable, indicating that GEMC1 is not essential for DNA replication (71).

In summary, although DUE-B interacts with key replication initiation proteins and can influence origin firing in certain experimental contexts, our combined zebrafish and *Xenopus* data indicate that it is not essential for vertebrate DNA replication or development. Determining the specific cellular contexts in which DUE-B becomes functionally important for DNA replication will be an important direction for future research.

## Materials and methods

### Animal care

Zebrafish were housed and cared for in strict accordance with protocols approved by the OMRF Institutional Animal Care and Use Committee. Adult fish were housed in an aquatic facility in tanks at a density of 10 fish per liter and maintained at a constant 26.5°C with 10-hour light and 14-hour dark cycles.

Eggs from adult lab bred female *Xenopus laevis* frogs (Xenopus 1, Cat# 4800) were used to prepare egg extracts. Testes from adult lab bred male *Xenopus laevis* frogs (Xenopus 1, Cat# 4293) were used to prepare sperm chromatin. All animals were healthy, not subjected to previous procedures, and no animal husbandry was performed. Frogs were housed at the Boswell Small Amphibians Research Facility at Stanford School of Medicine in compliance with Institute Animal Care and Use Committee (IACUC) regulations and all experiments involving frogs were approved by the Stanford University APLAC (Administrative Panel on Laboratory Animal Care).

### Bacterial cell lines

DH5α (New England Biorabs (NEB), Cat# C2987H), and Rosetta (DE3) pLysS (Novagen, Cat# 70956-4) bacteria were cultured in LB broth (Thermo Fisher Scientific, Cat# BP9723) at 37ᵒC for plasmid production and protein overexpression.

### IVTT system

IVTT reactions were performed using TnT SP6 High-Yield Wheat Germ Protein Expression System (Promega, Cat# L3261) according to the manufacturer’s instructions. Proteins were not purified after their production by IVTT. The IVTT reaction was directly added replication reactions as the source of a given protein. The optimal concentration of the template plasmid was titrated for each protein.

### TALEN production and microinjection

The TALEN expression constructs targeting the third exon of the *due-b (dtd1)* gene were assembled in pCS2TAL3-DD and pCS2TAL3-RR using Golden Gate Assembly as in (72). pCS2TAL3-DD and pCS2TAL3-RR were gifts from David Grunwald (Addgene plasmids # 37275 and 37276; RRID:Addgene_37275; RRID:Addgene_37276). mRNA encoding the TALE nuclease subunits was synthesized using the mMESSAGE mMACHINESP6 Transcription Kit (AM1340; ThermoFisher Scientific) and purified using RNeasy Mini Kit (74104; Qiagen).

125pg of each TALEN mRNA was injected into 1-cell stage zebrafish embryos. TALEN-injected embryos were raised to adulthood and assessed for transmission of insertions/deletions at the TALEN cut site by Pst1 restriction digest of PCR amplicons. F1 fish from mutation transmitting F0 animals were raised to adulthood and analyzed for mutation by PCR (*Forward primer*: tgtaatacgactcactatagggCAGGTTTTTCATCCCTGCAT; Reverse Primer: aaaacgacggccagtCTGCTCCAGCATGTTGTTGT) and restriction digest. The specific *due-b* mutation was identified by Sanger sequencing.

### Genotyping

Wild-type, heterozygous, and *due-b^omf202/omf202^* genotypes were identified by PCR analysis using the following primers: CS929: CTGGAGCAGCTTAGAGAAACCT; CS930: TGTGTGAGTCAGTTCACTCTG; CS931: GTGTGAGTCAGTTCACTCCT; CS791: GCGTTTAACAACAGTAGGCAATCA. All primers were purchased from Integrated DNA Technologies.

### *in situ* hybridization

*In situ* hybridization protocol was followed as in (73). Briefly, ISH probes were constructed by PCR from genomic DNA and isothermal assembly into HindIII-BamHI digested pUC57-Amp plasmids flanked byT3 andT7 promoters for sense (coding RNA) and anti-sense strand synthesis, respectively. Primers used to amplify probe sequences are as follows: *due-b* forward: accctcactaaagggaaGAGCGAGCGTAACAGTTGGA; *due-b* reverse: acgactcactatagggcTGGCGCTGGGATCTACTTTT. Dioxygenin (DIG)-labelled RNA probes were synthesized with T3 and T7 polymerase with reagents from Roche. Embryos were manually dechorionated with forceps, then fixed in 4% paraformaldehyde in 1X PBS overnight. Fixed embryos were dehydrated in 100% methanol and stored at −20°C. Embryos were proteinase K treated for 10 minutes. The protocol was programmed into an Intavis VSi machine to automate the prehybridization and washing procedures. Embryos were removed from the machine and incubated in DIG-labelled RNA probes overnight in a 70°C water bath.

Embryos were then returned to the machine for washing, pre-incubation with Bovine Serum Albumin, incubation with anti-DIG antibody, and additional washing. Labelled embryos were imaged with a Nikon SMZ1500 stereomicroscope with Andor Zyla CMOS camera.

### Quantitative RT-PCR

RNA was isolated from 50-80 embryos using Tri-Reagent (Invitrogen AM9738) following protocol from (74). mRNA was converted to cDNA using the AccuScript High Fidelity Reverse Transcriptase (600089; Agilent, Santa Clara, CA). qPCR was done on a Roche LightCycler 480 with the SYBR Green I Master (04707516001; Roche) master mix. Three technical replicates were performed for each sample. The relative mRNA expression was calculated using the 2-ΔΔ^Cq^ method. The following primer sequences were used: *due-b* forward: CGGGCAGTTTGGAGCAAAAA; *due-b* reverse: TGGACAGCAGTTTGGGATCT; r*pl13a*: TCTGGAGGACTGTAAGAGGTATGC; *Rpl13a* reverse AGACGCACAATCTTGAGAGCAG.

### Western blot

Zebrafish experiments: Bud stage embryos (10 hpf) were collected, dechorionated with pronase (Roche 11459643001) deyolked, and lysed in RIPA buffer (150mM NaCl, 50mM Tris pH8, 1mM EDTA, 0.5% deoxycholate, 1%NP-40, 0.1%SDS). 50ug of total protein were loaded onto SDS-PAGE gel. Anti-human DUE-B antibody was a gift of Michael Leffack. The B-Actin Antibody was purchased from Abcam (ab6276).

Xenopus experiments: All samples were boiled in 1x Laemmli buffer and run on 4-15% Mini-PROTEAN TGX Precast Protein Gels (Bio-Rad #4561086) using Tris-Glycine-SDS Running Buffer (Boston BioProducts #BP-150). EZ-Run Prestained Rec Protein Ladder (Fisher Scientific #BP36031) or Precision Plus Protein Dual Color Standard (Bio-Rad #1610374EDU) was run alongside samples to infer protein band sizes. For immunoblotting, proteins were transferred from the gel onto a PVDF membrane (VWR #10061-492) in transfer buffer (Fisher Scientific #NC9297917). Membranes were blocked in 1x PBST (1x Phosphate Buffered Saline (PBS) (Fisher Scientific, Cat# NC9140736), 0.05% Tween 20) containing 5% (w/v) nonfat milk for 1 hour at room temperature with gentle shaking. Following a wash in 1xPBST, the membrane was incubated with primary antibodies diluted in 1x PBST containing 1% (w/v) BSA overnight at 4ᵒC with gentle shaking. Membranes were washed three times with 1x PBST and incubated with secondary antibody diluted in 1x PBST + 5% nonfat milk for 1 hour at room temperature with gentle shaking. Membranes were washed three times with 1x PBST, developed using SuperSignal West Pico PLUS Chemiluminescent Substrate (Fisher Scientific, Cat# PI34577), and imaged using Azure c600 (Azure Biosystems).

### Cloning

The *Xenopus laevis dtd1* gene was cloned as follows. The *dtd1* gene was PCR amplified from *Xenopus laevis* cDNA using oLAS045 and oLAS046. The resulting PCR product was then cloned into the NotI-HF digested pFastBac1 vector using NEBuilder HiFi mix, resulting in the plasmid [pLAS023]pFastBac-xDUEB. The *dtd1* gene cloned from cDNA prepared in our lab was identical to the *dtd1* sequence NM001093157.1 from the NCBI database. The *dtd1* gene from our cDNA was used for all subsequent cloning and experiments.

To express *Xenopus laevis* DUE-B with a His6 tag at the C-terminus, the *dtd1* gene was PCR amplified from pLAS023 using oLAS049 and oLAS050. The backbone was generated by PCT amplifying pET-28b(+) using oLS019 and oLS023. The two PCR products were assembled using NEBuilder HiFi mix, yielding the plasmid [pLAS026]pET28-xDUEB-His6.

To express untagged DUE-B in IVTT, the *dtd1* gene was PCR amplified from pLAS023 using oLAS060 and oLAS063. The resulting untagged *dtd1* gene product was then digested with SalI and SacI and cloned into the SalI-SacI region of pSP64 Poly(A), resulting in the plasmid [pLAS031]pSP64-xDUEB.

To express DUE-B^FLAG^ in IVTT, the *dtd1* gene was PCR amplified from pLAS023 using oLAS060 and oLAS061. The resulting untagged *dtd1* gene product was again PCR amplified using oLAS060 and oLAS062 to add the C-terminus FLAG tag. The resulting *dtd1-FLAG* gene product was then digested with SalI and SacI and cloned into the SalI-SacI region of pSP64 Poly(A), resulting in the plasmid [pLAS030]pSP64-xDUEB-FLAG.

To express ^FLAG^DUE-B in IVTT, the *dtd1* gene was PCR amplified from pLAS023 using oLAS063 and oLAS064. The resulting untagged *dtd1* gene product was again PCR amplified using oLAS063 and oLAS065 to add the N-terminus FLAG tag. The resulting *FLAG-dtd1* gene product was then digested with SalI and SacI and cloned into the SalI-SacI region of pSP64 Poly(A), resulting in the plasmid [pLAS032]pSP64-FLAG-xDUEB.

### Protein purification

The native purification of antigens was performed as follows. The Rosetta (DE3) pLysS cell pellet overexpressing the His6-tagged antigen was thawed in Lysis buffer containing 1 mM PMSF, and sonicated. The lysate was cleared by ultracentrifugation at 35,000 rpm for 40 min at 4°C in a Beckman Type 70 Ti rotor. Cleared lysate was incubated with 1 mL of Ni-NTA resin for 1 hr at 4°C. The protein was eluted with 400 mM imidazole in Lysis buffer containing 1 mM PMSF.

The denaturing purification of antigens was performed as follows. The Rosetta (DE3) pLysS cell pellet overexpressing His6-tagged antigen was thawed in Denaturing Lysis buffer (50 mM HEPES pH7.5, 500 mM NaCl, 10 mM imidazole, 8 M Urea) containing 1 mM PMSF, and sonicated. The lysate was cleared by ultracentrifugation at 35,000 rpm for 40 min at room temperature in a Beckman Type 70 Ti rotor. Cleared lysate was incubated with 1 mL of Ni-NTA resin for 1 hr at room temperature. The protein was eluted with 400 mM imidazole in denaturing elution buffer (50 mM HEPES pH7.5, 500 mM NaCl, 6 M Urea, 1 mM PMSF).

### Cell cycle analysis

EdU experiments were based on Sansam et al. 2010 (5). Briefly, after fertilization, individual embryos were placed in single wells of 96-well plate. At 24hpf, embryos were dechorionated with pronase (Roche 11459643001), gently washed in E3 fish water(5mM NaCl, 0.17mM KCl, 0.33mM CaCl2, 0.33mM MgSO4). They were labelled with 20mM 5-ethynyl-2’-deoxyuridine (EdU, ThermoFisher A10044) in DMSO for 20 minutes, transferred to fresh E3 medium and allowed incubated for an additional 15 min at 28.5°C. Embryos were then washed thoroughly in 1XPBS and placed in ice. Embryos were disaggregated with a multi-channel pipet into a single cell suspension and a small portion of cells were removed for genotyping PCR. The remaining cells were fixed in 70% ethanol overnight. Cell suspensions from the same genotype were pooled and prepared for flow-cytometry with the click-chemistry. Flow cytometry data was quantified by FlowJo (TreeStar, Inc.)

### Tail fin regeneration

Tail fin regeneration assays were performed as in described in (60).

### Xenopus egg extract preparation

*Xenopus laevis* egg extracts were prepared following published protocols (75). Frogs were primed with 75 IU of human chorionic gonadotropin (CHORULON) (Merck, Cat# 133754) 5-7 days before extract preparation. To induce ovulation, 660 IU of CHORULON was injected 20-21 hr before extract preparation. High-speed supernatant (HSS) was prepared from 5-6 adult female frogs, and nucleoplasmic extract (NPE) was prepared from 15-20 frogs. Sperm chromatin was purified from adult male frog testes and used for NPE preparation.

### Ensemble DNA replication assay

To validate antibodies and recombinant proteins, ensemble DNA replication experiments were performed following a detailed protocol described in a recent methods article (67). Since DUE-B is dispensable for Mcm2-7 double-hexamer loading, all licensing reactions were performed with HSS depleted of DUE-B.

### Antibody Preparation

Custom polyclonal antibodies were generated as previously described (67). Briefly, the His6 tagged antigen sequence was cloned into pET28b(+). The antigen was expressed in Rosetta (DE3) pLysS (Novagen, Cat# 70956-4) upon induction with 0.5-1.0 mM IPTG for 3 hrs at 37°C or overnight at 16°C. Soluble antigens were purified in native buffer, whereas insoluble antigens were purified in denaturing buffer. In most cases, the Ni-NTA eluate (still containing 400 mM imidazole) was used to immunize rabbits. For some soluble antigens, additional chromatography was used to polish the protein preparation. All antibodies used in this study were raised by Cocalico Biologicals.

All antibodies used for *Xenopus* egg extract experiments were affinity purified. 1-5 mg of purified antigen was cross-linked to 1 mL of AminoLink Plus Coupling Resin (Fisher Scientific, Cat# PI20501) according to the manufacturer’s protocol. 10-30 mL of serum was incubated with the affinity resin for 30 minutes at room temperature. The resin was washed twice with 10 mL of 1x PBS + 500 mM NaCl, and twice again with 10 mL of 1x PBS. Next, antibodies bound to the affinity resin were eluted 5-6 times with 1.0 mL of 200 mM glycine-HCl pH2.5. Each elution was collected into an tube containing 0.1 mL of 1M Tris-HCl pH9.0, which immediately neutralized the glycine. Eluted fractions were pooled and dialyzed against 1 L of 1x TBS (50 mM Tris-HCl pH8.0, 150 mM NaCl) + 10% (w/v) sucrose. The IgG concentration was measured using a Nanodrop, normalized to 1 mg/mL via dilution or spin-concentration, aliquots were flash frozen in liquid nitrogen, and stored at −80°C.

### Data availability

Flow cytometry data will be made available in a public repository upon publication.

## Acknowledgements

We thank Dr. Michael Leffak for the generous gift of serum raised against human DUE-B protein. We thank the Flow Cytometry Core Facility at the Oklahoma Medical Research Foundation for their assistance. We thank Joseph Siefert for providing assistance with the *in situ* hybridization. We thank Dr. Susannah Rankin for helpful discussions. We thank the members of the Chistol lab for feedback. This work was supported by a NIGMS R01 grant to C.L.S. (1R01GM121703), a NSF CAREER award to G.C. (2144481), and a NIGMS R35 grant to G.C. (GM147060). L.A.S. was supported in part by the Blavatnik Family Fellowship Fund.

## Author contributions

CLS, LAS, EAM, GC, and CGS conceived and designed experiments. CLS designed the *due-b* TALENs and generated the mutant. EAM and CGS performed cohort analyses. DG assisted with fish husbandry and genotyping. LAS performed the *Xenopus* egg extract experiments. CGS, CLS, and GC wrote the manuscript.

